# ARGus: A Co-assembly workflow for MAG generation, ARG detection, and virulence analysis

**DOI:** 10.64898/2026.05.22.727233

**Authors:** Nivedita Puliadi Subramanian, Dennis D. Krutkin, Scott T. Kelley

**Author notes:** Corresponding Author. Department of Biology, San Diego State University, San Diego, California 92182, USA.

## Abstract

The emergence of antibiotic resistance among pathogenic bacteria is a significant global health challenge with multidrug resistance becoming increasingly common. Moreover, since antibiotic resistance genes (ARGs) can be transferred horizontally more bacteria are rapidly evolving resistance. In addition, emerging bacterial pathogens continue to arise from a combination of urbanization, animal agriculture, global movements of people, and inadequate sewage infrastructure. Researchers have begun applying deep sequencing and shotgun metagenomics to detect known and unknown pathogenic organisms and ARGs directly from environmental samples. Here, we describe a bioinformatics workflow that uses a co-assembly approach to assemble contigs across metagenomes and bin them into high coverage metagenomic assembled genomes (MAGs), while segregating out unbinned contigs that includes mobile elements (e.g., plasmids). The workflow includes annotation of coding sequences and differential determination of ARGs and virulence factors (VF) within the sets of both MAG genome bins and unbinned contigs and allows quantification of MAG, ARG and VF abundances for ecological (alpha and beta diversity) and network analyses. Workflow analysis of metagenomic samples collected from the heavily polluted Tijuana River identified hundreds of MAGs, including many high-quality bins and many novel potential pathogens, and found the vast majority of ARG sequence matches in the unbinned contigs. A combined network analysis found strong correlations (r > 0.90) between ARGs and specific MAGs, indicating which bacterial species is likely to contain the ARG. This workflow provides a powerful approach for public health metagenomics studies of emerging pathogens and ARGs.

## Introduction

The emergence of antibiotic resistance among pathogenic bacteria has become a significant global health challenge (Bukari et al., 2025; Frieri et al., 2017; Voicu & Ahmet, 2021). By 2050, it is estimated that deaths related to infectious diseases could reach as many as 10 million per year (De Nies et al., 2021). Because the exact causative microorganisms of infections are often unknown, patients are frequently treated with broad-spectrum antibiotics. The improper or unregulated use of antibiotics, including self-medication without prescriptions, has substantially contributed to the rise and spread of antibiotic resistance (Frieri et al., 2017).

Antibiotic resistance works via four primary mechanisms: enzymatic inactivation, efflux pumps, target alteration, and reduced permeability. In enzymatic inactivation, bacteria produce enzymes that hydrolyze or chemically modify antibiotics, rendering them inactive. Efflux pumps are proteins embedded in the bacterial cell membrane, such as ABC transporters, that actively expel antibiotics, reducing their intracellular concentration and effectiveness. Target alteration occurs when mutations or chemical modifications change the antibiotic’s binding site, preventing effective drug interaction. Finally, reduced cell permeability resulting from alterations in membrane proteins limits antibiotic uptake and restricts antibiotics from reaching their intracellular targets (Álvarez-Martínez et al., 2020; MacGowan & Macnaughton, 2017).

Antibiotic resistance traits can arise in two primary ways. One way is via spontaneous mutations in the bacterial genome, which can lead to traits such as reduced permeability and target alterations (Belay et al., 2024). The other way is via horizontal gene transfer (HGT). Mobile genetic elements (MGEs) such as plasmids, transposons, and integrons, facilitate the transfer of antibiotic resistance genes (ARGs), mainly efflux pumps and inactivation enzymes, between bacteria via HGT (Belay et al., 2024; Fuentes et al., 2019). HGT enables the acquisition of resistance genes from other bacteria and happens through (1) conjugation, the direct transfer of genetic material between bacterial cells through physical contact, often mediated by plasmids; (2) transduction, the transfer of genetic material between bacteria via bacteriophages; and (3) transformation, the uptake of free DNA fragments from the surrounding environment (Álvarez-Martínez et al., 2020; Shaikh et al., 2015).

The Tijuana River is an environmental hotspot for pathogens carrying ARGs, which can cause illness after a single exposure. These pathogens flow through the Tijuana River Estuary into the Pacific Ocean, contributing to the spread of antibiotic-resistant bacteria and raising public health concerns for individuals in contact with the water (Fuentes et al., 2019; Pendergraft et al., 2023; Rico et al., 2024). Pathogenic microorganisms cause disease by invading, colonizing, and damaging the host. Pathogenicity is often driven by traits encoded by virulence factors (VFs), and the presence of antimicrobial resistance (AMR) can exacerbate disease severity. Opportunistic pathogens carrying both VFs and ARGs are more likely to persist and proliferate, increasing the risk of multidrug-resistant infections (De Nies et al., 2021).

Traditional methods, such as culture-based assays and quantitative PCR (qPCR), are limited in their ability to detect novel bacteria or ARGs, particularly those from uncultivable or previously uncharacterized bacteria. In contrast, shotgun metagenomics enables detection of both known and novel ARGs, as well as potential new pathogenic species. In metagenomic studies, ARG mobility is often inferred by identifying their presence on mobile genetic elements (MGEs) within the unbinned fraction of co-assembled contigs. While traditional culturing and PCR can monitor ARGs, they cannot reliably determine which ARGs reside within specific genetic elements, such as chromosomes or MGEs. However, a shotgun metagenomic approach that sequences DNA fragments isolated directly from the microbial communities can be used to probe both taxonomic diversity and ARG diversity in complex microbial environments without the need for culturing or prior knowledge of ARG sequences (B. Li et al., 2022).

Use of k-mer methods, such as Kraken and Kaiju, can efficiently detect the presence of antibiotic resistance genes (ARGs) or virulence factors (VFs); however, they cannot reliably determine whether ARGs/VFs reside on chromosomes or plasmids/mobile genetic elements. Additionally, these methods rely on DNA k-mers (Kraken) or protein-level k-mers (Kaiju), making it challenging to analyze the underlying DNA sequence (Menzel et al., 2016; Wood & Salzberg, 2014).

Here, we present a workflow based on metagenome co-assembly that enables the reconstruction of both genomes and mobile genetic elements. This approach improves strain-level genomic resolution by co-assembling metagenomes from multiples samples and binning output contigs, resulting in more complete assemblies compared to read-based or individual assembly methods. Our work also collects unbinned contigs that may include extra-chromosomal or mobile elements (e.g., plasmids) and analyzes both MAG binned and unbinned contigs for the presence of ARGs. The workflow also calculates the relative abundances of MAGs and ARGs, annotates contigs and determines potential MAG pathogenicity by identifying the types and number of virulence factors contained in each MAG. We tested our workflow using metagenomes collected from two different studies of the heavily contaminated Tijuana River. Our results demonstrate that co-assembly enhances the detection of ARG diversity and location (genomes vs. mobile elements), identifies virulence factors along with their genomic contexts, and uncovers novel correlations between ARGs and MAGs that would be missed by short-read classification methods alone.

## Materials and methods

### Data collection and computational resources

Raw metagenomic sequencing data for this study were obtained from two previously published studies of water samples collected from the United States portion of the Tijuana (TJ) River (Allsing et al., 2022; Shahar et al., 2024a). The TJ River flows south to north from Mexico into the United States, with approximately 25% of its length in the US, and deposits into the ocean estuary south of Imperial Beach, CA. The South Bay International Wastewater Treatment Plant (SBIWTP) plant was built north of Tijuana’s main wastewater pumping station to add secondary treatment but is inadequate to handle the waste volume. Both treated and raw sewage is released into the river which cause extensive beach closures (Shahar et al. 2022). Allsing et al. generated 22 metagenomes from contaminated water samples collected from 6 sites on the United States portion of the river from November 2019 to February 2020. Shahar et al. generated 13 samples collected from two sites, one near the border at San Ysidro, CA and one near the estuary near the mouth of the river near Imperial Beach, CA at 7 different days from August 2020 to May 2021. Sample collection molecular method approaches were the same in both these studies. Briefly, water samples were collected using a telescopic dipper sampler were transferred to sterile Whirl-Pak bags and transported to the lab on ice. At the lab, water samples were immediately filtered through 0.22 μm Sterivex filters and between 50 and 100 mL of water was obtained per sample. DNA was extracted directly from the filtered water samples using a Qiagen DNeasy PowerWater Sterivex kit. Purified DNA was sent to the Microbial Genome Sequencing Center (MiGS; Pittsburgh, PA, USA), for Nextera XT DNA library constructions and sequencing on a NextSeq200 platform at a depth of approximately 25M. The Allsing et al. data were downloaded European Nucleotide Archive (ENA), project accession number PRJEB57859 and the Shahar et al. data are available at NCBI Sequence Read Archive (SRA), Bioproject PRJNA1423341.

Bioinformatic analyses were performed on two high-performance servers. The first server was equipped with a single Intel Xeon W-2295 CPU (18 cores, 36 threads, 3.0 GHz) and 250 GiB of RAM. The second server had two Intel Xeon Silver 4416+ CPUs (20 cores per CPU, 40 cores total, 40 threads total) and 1.0 TiB of RAM. Both servers ran a 64-bit Linux operating system.

### Sequence preprocessing, co-assembly, and MAGs recovery

Paired-end raw reads were preprocessed using an all-in-one tool named fastp v0.23.2 to remove adapter contamination and polyG tails, trim low-quality bases, and filter short reads. Quality control reports were generated per sample in JSON and HTML formats, and the results were summarized using fastp_summarize.py to produce the overall statistics before and after filtering (S. Chen et al., 2018). Fastq-pair v1.0 was used to ensure that each read in one file had a corresponding mate in the other (Edwards & Edwards, 2019). Next, the paired-end reads were co-assembled from all the metagenomic samples, which included a variety of closely related strains within the same species (Delgado & Andersson, 2022). This was performed using MEGAHIT v1.2.9, the first Next-Generation Sequencing (NGS) metagenome assembler (D. Li et al., 2016). The Metagenome-assembled genomes (MAGs) were recovered from co-assembled contigs using MaxBin 2.0 v2.2.7, based on tetranucleotide frequencies and read coverage levels, where each bin represents a single genome (Wu et al., 2014, 2016). The taxonomy of each bin was assigned with the help of 120 single-copy bacterial and 53 archaeal marker genes using the Genome Taxonomy Database Toolkit (GTDB-Tk v2.4.0) (Chaumeil et al., 2020, 2022).

### Gene prediction and virulence factor identification

The coding sequences (CDSs) of each binned genome were predicted and translated using Prodigal v2.6.3 (Hyatt et al., 2010). Virulence factor protein sequences (Set B) were downloaded from the Virulence Factor Database (VFDB; accessed April 2025, https://www.mgc.ac.cn/VFs/download.htm) (L. Chen et al., 2016). A protein BLAST database was created from these sequences using makeblastdb (NCBI BLAST+ v2.16.0) (Camacho et al., 2009). A Python script (blastp_for_VFs.py) was used to align the CDSs of each binned genome against this database using BLASTp with 100% query coverage, an E-value cutoff of 10, a maximum of five hits per query, and output format 5 (XML). Hits to virulence factors were exported in a .txt file containing the following information: Bin_ID, Query.ID, Query_Length, Hit.ID, Hit_Description, Subject_Length, Score, Bits, E-value, Identities, Positives, Gaps, Query_Start_Position, Query_End_Position, Subject_Start_Position, Subject_End_Position, Query and Subject Alignment, as well as the complete GenBank record of each hit. Another Python script (Top_VF_hits.py) was then used to extract the top hit for every contig within each MAG, and a virulence factor count table was subsequently generated for each MAG across the main virulence factor categories.

### Abundance Estimation of Metagenome-Assembled Genomes

The relative abundance of the binned genomes was calculated by first creating an index from the co-assembly contigs, and the reads were then aligned against this index using Bowtie2 v2.5.4 (Langmead & Salzberg, 2012). The resulting .sam files were converted to .bam files and sorted using SAMtools v1.22.1 (Danecek et al., 2021). The relative abundance of the MAGs was subsequently calculated from the sorted .bam files using CoverM v0.7.0 (Aroney et al., 2025). A Python script (MAG_relative_abundance.py) was used to filter out unmapped entries, retain only bacterial MAGs, calculate total relative abundance across samples, generate a feature table with samples as rows and MAGs as columns, and rename MAG columns using the lowest resolved GTDB-Tk taxonomy combined with a standardized MAG ID. The final processed table was saved as a .csv file for downstream analyses.

### Virulence Factor Analysis

Heatmaps of virulence factor (VF) counts per VF category per MAG were generated using the Python script VF_count_taxonomy.py. The VF count table per MAG was merged with the taxonomic classification table obtained from GTDB-Tk v2.4.0, removed entries without valid taxonomy information, and assigned each MAG a unique standardized identifier. The lowest resolved taxonomic rank was extracted from the GTDB-Tk classification and combined with MAG ID to create informative row labels for visualization. VF categories were renamed using acronyms and ordered consistently across samples. Total VF counts per MAG were calculated, and MAGs were sorted in descending order based on total VF abundance. The script produced two heatmaps: one including all MAGs and another displaying the top 20 MAGs with the highest VF counts. Higher color intensity represents higher VF abundance within each VF category.

### Processing of Unbinned contigs for ARG detection

A bash script (segregate_unbinned_contigs.sh) was used to identify all the unbinned contigs by comparing the genome-assigned contigs with the co-assembled contigs. Any co-assembled contigs not assigned to genome bins were segregated as unbinned. The sequences of the unbinned contigs were then extracted using seqtk v1.4 (H. Li, 2012), and their coding sequences (CDSs) were predicted and translated using Prodigal v2.6.3 (Hyatt et al., 2010). Antibiotic inactivation and antibiotic efflux genes were downloaded from the Comprehensive Antibiotic Resistance Database (CARD) using the qualifiers *part_of*, *is_a*, *participates_in*, *has_part*, and *nucleotide* (Alcock et al., 2023a). A protein BLAST database was created from these sequences using makeblastdb (NCBI BLAST+ v2.16.0) (Camacho et al., 2009).

### ARG Identification and Quantification in Unbinned contigs

A python script (blastp_for_ARGs_CDS.py) was used to align the CDSs of the unbinned contigs against this database using BLASTp with 100% query coverage, an E-value cutoff of 10, a maximum of five hits per query, and output format 5 (XML). Hits to both inactivation and efflux ARGs were exported in a .txt file separately containing the following information: Bin_ID, Query.ID, Query_Length, Hit.ID, Hit_Description, Subject_Length, Score, Bits, E-value, Identities, Positives, Gaps, Query_Start_Position, Query_End_Position, Subject_Start_Position, Subject_End_Position, Query and Subject Alignment, as well as the complete GenBank record of each hit when available. The script also extracted the top protein hit for each unbinned contig from the BLASTp XML output. For each top hit, the corresponding protein sequence was retrieved from NCBI and saved as a .faa file. The linked nucleotide sequence (CDS) was also fetched from the NCBI nuccore (Sayers et al., 2012), when available and saved as a .fna file. Duplicate headers in the .fna file were removed using seqkit v2.8.2. The cleaned .fna file was then indexed, and the reads were aligned using Bowtie2 v2.5.4 (Langmead & Salzberg, 2012). The resulting .sam files were converted to .bam files and sorted using SAMtools v1.22.1 (Danecek et al., 2021). Finally, the read counts of the ARGs from unbinned contigs were calculated from the sorted .bam files using CoverM v0.7.0 (Aroney et al., 2025).

### Protein-CDS Mapping for Unbinned contigs

Another Python script (protein_CDS_mapping.py) was used to extract the top protein hit for each unbinned contig from the BLASTp XML output. For each hit, the corresponding protein sequence was retrieved from NCBI and saved as a .faa file. The script also retrieved the linked CDS nucleotide sequence from the NCBI nuccore database, when available, and saved it as .fna file. For each CDS, additional information including gene name, locus tag, and CDS ID were recorded. A mapping table was generated to link previously unknown ARG names and descriptions, connecting contig IDs to their CARD protein accessions, CARD protein descriptions, NCBI nucleotide IDs, gene names, locus tags, and CDS IDs.

### Integration of ARG Read counts with Protein-CDS Mapping

A subsequent Python script (merging_final_feature_table.py) integrated the ARG read count data with the protein-CDS mapping information. The script merged the read count table of unbinned contigs with the protein-CDS mapping table using contig IDs in the feature table and the CDS IDs in the mapping table as the merge keys. Only the read counts and the corresponding CARD protein accessions, CARD protein descriptions, NCBI nucleotide IDs, and CDS names were retained. The resulting combined table was saved as a .tsv file for downstream analyses.

### Final ARG Table Processing and Summarization

Duplicate entries based on CARD protein accessions were first removed separately for inactivation and efflux CDS read count tables using a python script (Duplicates_removal_and_merging.py). The cleaned tables were then merged to create a comprehensive unbinned ARG dataset. For each ARG, total read counts across all samples were calculated, and the top 10 ARGs by total read count were summarized. The combined table was transposed so that samples became rows and ARGs became columns, rows containing only zero values were removed, and ARGs were renamed based on CARD protein descriptions. The final processed table was saved as a .csv file for downstream analyses.

### Analysis Pipeline

The entire workflow described in Figure 1 has been configured as the ARGus Snakemake workflow. The complete details of the Snakemake pipeline, including a list of software and databases used, can be found at https://github.com/ddkrutkin/ARGus/tree/main.

**Figure 1.**
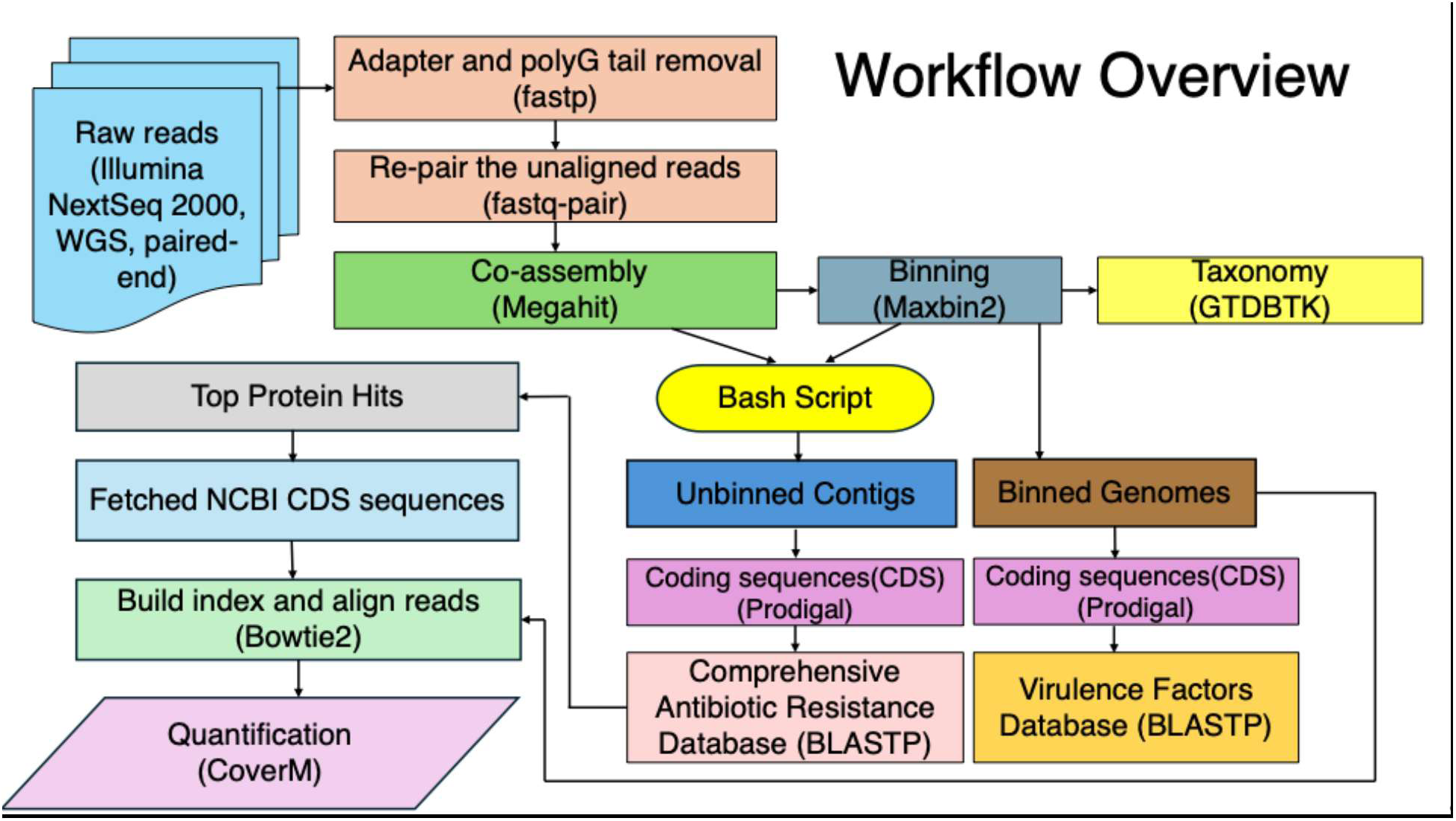
Overview of the ARGus pipeline for metagenomic processing and analysis. Raw paired-end reads from Illumina NextSeq 2000 WGS were preprocessed to remove adapters and polyG tails using fastp v0.23.2 and re-paired with fastq-pair v1.0. Cleaned reads were co-assembled with MEGAHIT v1.2.9 to generate contigs, which were binned into MAGs using MaxBin 2.0 v2.2.7 and taxonomically classified with GTDB-Tk v2.4.0. Unbinned contigs were separated from MAGs for downstream analyses. Coding sequences (CDSs) were predicted from both MAGs and unbinned contigs using Prodigal v2.6.3. Antibiotic resistance genes (ARGs) were identified by aligning CDSs from unbinned contigs against a comprehensive antibiotic resistance database using BLASTP; top hits were extracted via a Python script, corresponding NCBI CDS sequences were retrieved, an index was constructed from these sequences, reads were aligned against this index using Bowtie2 v2.5.4, and read counts were quantified with CoverM v0.7.0. Virulence factors (VFs) were identified similarly from MAGs using the Virulence Factor Database (VFDB). For each MAG, top hits were extracted via a Python script, and a VF count table was generated across the main VF categories. Relative abundances of MAGs were calculated by aligning MAGs against the assembled contigs using Bowtie2 v2.5.4 and quantified with CoverM v0.7.0.

### Statistical analysis

Patterns of microbial community composition were explored across samples using Non-metric Multidimensional Scaling (NMDS). The relative abundances of MAGs and read counts of ARGs were subjected to robust centered log-ratio (rCLR) transformation, and Euclidean distances were calculated in R prior to NMDS and visualization using the vegan package (Oksanen et al., 2022). PERMANOVA tests were also performed using vegan with 9999 permutations. The rCLR-transformed MAG and ARG datasets were then combined, and the pybootnet program (Akhavan & Kelley, 2025) was used to perform bootstrap Spearman correlations and generate network graphs to identify significant correlations between ARGs and specific MAGs.

## Results

As an initial quality control step, sequencing reads from both datasets were assessed before and after filtering. For Dataset 1, the average GC content before filtering was 47.59% and after filtering was 47.54%, with approximately 99.25% of reads retained. For Dataset 2, the average GC content remained unchanged before and after filtering (46.94%), and approximately 99.99% of reads were retained following filtering. In Dataset 1, the average unbinned contig length was 443 bp, with a total unbinned length of 2,003,629,355 bp, whereas the total nucleotide length of all metagenome-assembled genomes (MAGs) was 797,868,231 bp, indicating that approximately 28.5% of the total sequence length was recovered in MAGs. In Dataset 2, the average unbinned contig length was 504 bp, with a cumulative unbinned length of 1,277,735,871 bp and a total MAG nucleotide length of 687,211,786 bp, corresponding to approximately 35% binned sequences.

MaxBin2 produced 274 prokaryotic bins in Dataset 1, of which 273 were bacterial and one was archaeal, classified at the genus level as *Nitrosarchaeum*. In Dataset 2, 315 prokaryotic bins were recovered, including nine archaeal bins, with the remaining bins classified as bacterial. Among the archaeal bins, three were classified as Marine Group II (MGII) archaea belonging to the phylum *Thermoplasmatota*, while the remaining six were unclassified. For Dataset 1, GC content across recovered bins ranged from 21.7% to 69.9%, and genome sizes spanned from small partial genomes (∼100 kb) to large genomes exceeding 11 Mb. Only a limited number of bins achieved complete (100%) or near complete (>90%) genome recovery, with the majority representing partial genomes of varying completeness. Dataset 2 showed a similar GC content range (approximately 22–70%), with genome sizes extending from highly fragmented assemblies (<200 kb) to large genomes exceeding 8 Mb.

### NMDS Analysis of MAG Relative Abundance Across Two Datasets

Figure 2 shows the NMDS plot for Dataset 1, samples from six sites and four collection dates generally clustered together, with samples 21, 10, 16, 2, 22, and 11 positioned farther from the main cluster, suggesting distinct MAG compositions in these samples. Permutational analysis of variance (PERMANOVA) analysis indicated that MAG composition differed significantly among sites (pseudo-F = 2.6; R² = 0.45; p = 0.003), but not among collection dates (pseudo-F = 0.86; R² = 0.13; p = 0.64). Figure 3 shows the NMDS plot for Dataset 2, which includes samples from two sites collected across seven dates. Samples from the same site generally clustered together. Specifically, samples 3, 5, 9, and 7 formed one cluster, while samples 6 and 10 formed a separate cluster, indicating intra-cluster similarity in MAG composition. PERMANOVA analysis indicated that MAG composition differed significantly among sites (pseudo-F = 4.6; R² = 0.29; p = 0.003), whereas differences among collection dates were not significant (pseudo-F = 0.58; R² = 0.05; p = 0.75).

**Figure 2.**
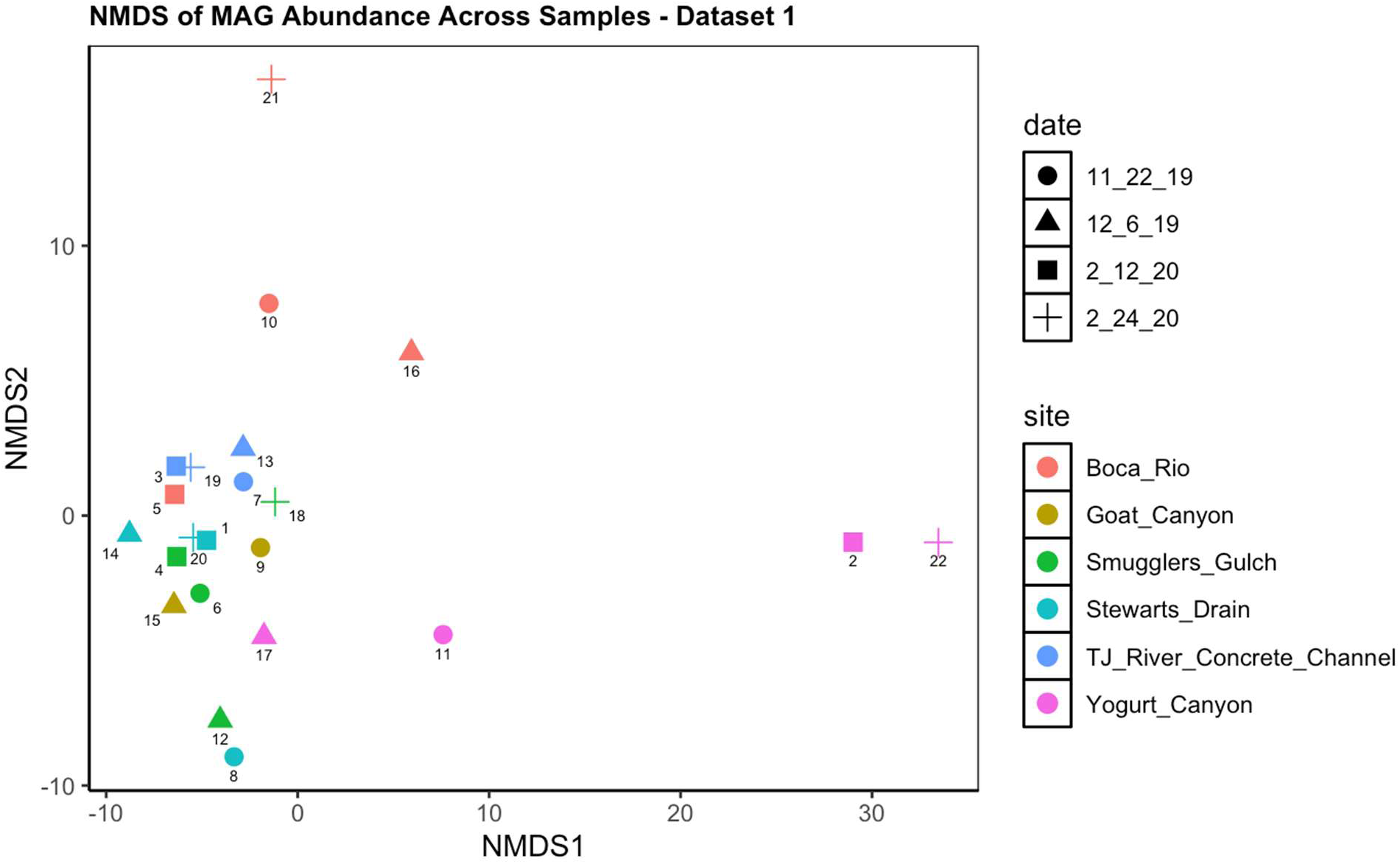
NMDS plot of MAG relative abundance across samples for Dataset 1. Relative abundances of MAGs were robust centered log-ratio (rCLR) transformed, and Euclidean distances were calculated to quantify dissimilarities between samples prior to NMDS. Each point represents an individual sample, with colors indicating sampling site and shapes indicating sampling date. Numbers next to points correspond to individual samples. NMDS stress = 0.065

**Figure 3.**
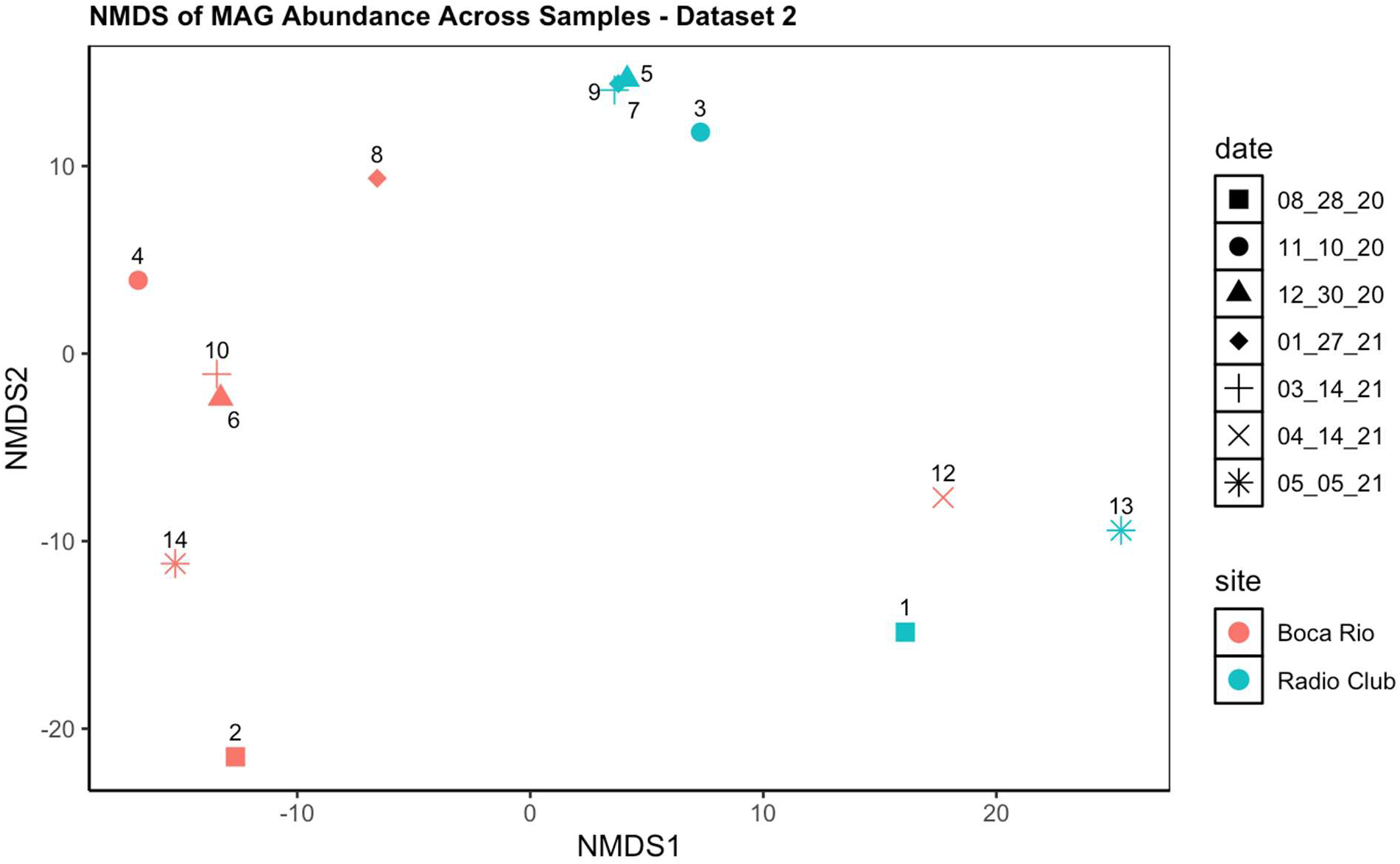
NMDS plot of MAG relative abundance across samples for Dataset 2. Relative abundances of MAGs were robust centered log-ratio (rCLR) transformed, and Euclidean distances were calculated to quantify dissimilarities between samples prior to NMDS. Each point represents an individual sample, with colors indicating sampling site and shapes indicating sampling date. Numbers next to points correspond to individual samples. NMDS stress = 0.058

### Distribution of Virulence Factors among Top 20 MAGs across Two Datasets

Figures 4 and 5 show heatmaps of virulence factors (VFs) for Datasets 1 and 2, respectively, presenting VF counts per VF category for the top 20 MAGs. Overall, the heatmaps reveal that most common VF categories include Adherence, Effector Delivery System, Immune Modulation, Motility, and Nutritional/Metabolic factors.

**Figure 4.**
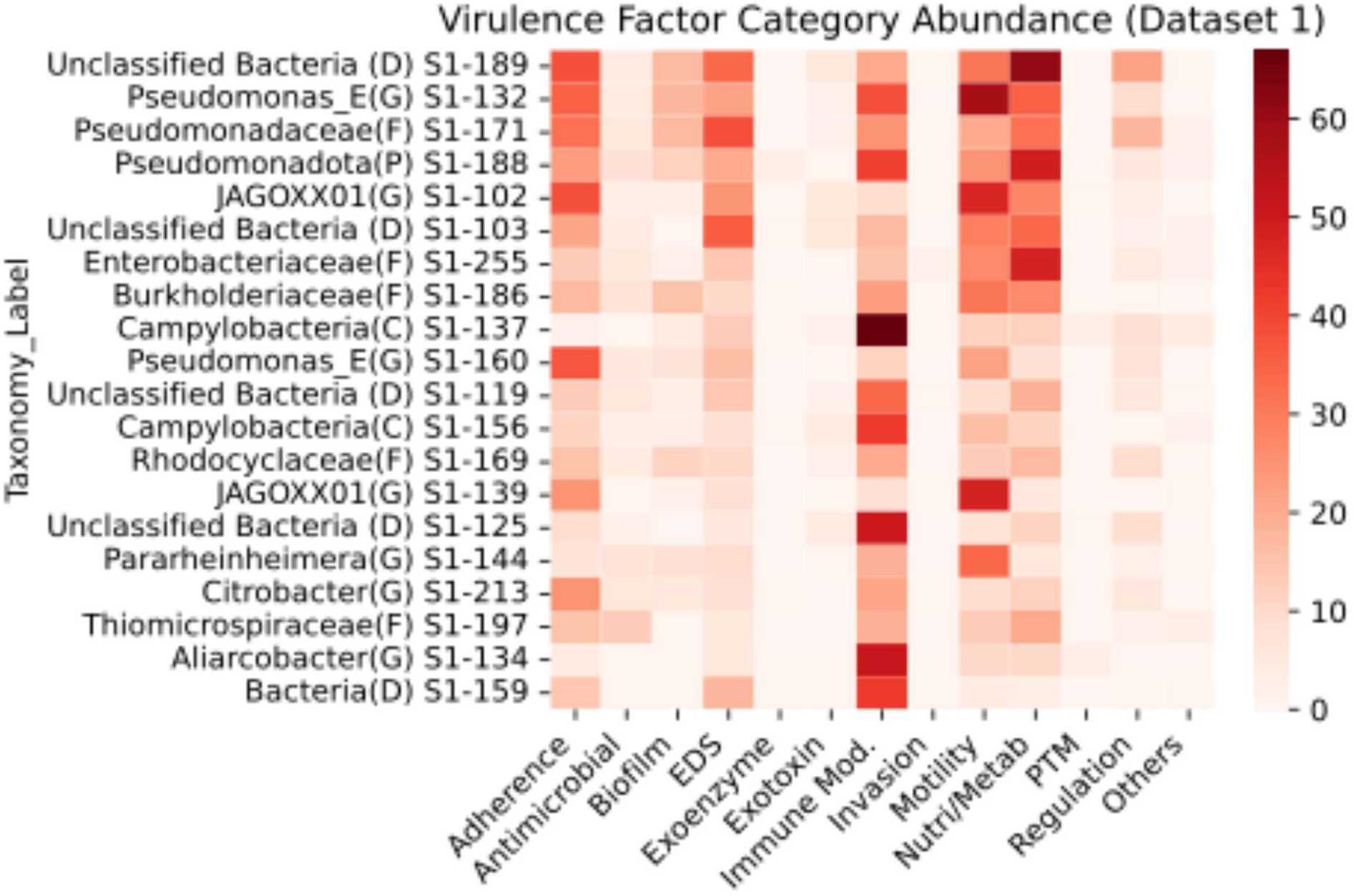
Heatmap showing virulence factor (VF) counts per VF category for the top 20 MAGs containing the most overall VFs in Dataset 1. VF categories on the X-axis were abbreviated (e.g., EDS = Effector Delivery System; Immune Mod. = Immune Modulation; Nutri/Metab = Nutritional/Metabolic Factor; PTM = Post-Translational Modification). The Y-axis lists MAGs by their most resolved taxonomic rank and numerical identifier, where S1 indicates the dataset to which the study belongs (Allsing et al., 2022), and the numbers following it indicate the bin number assigned by Maxbin2 v2.2.7. Color intensity reflects VF count, with darker colors indicating higher counts. Taxonomic abbreviations: D = Domain, P = Phylum, C = Class, F = Family, and G = Genus. Supplementary Table 1 contains the complete table for all MAGs in Dataset 1.

**Figure 5.**
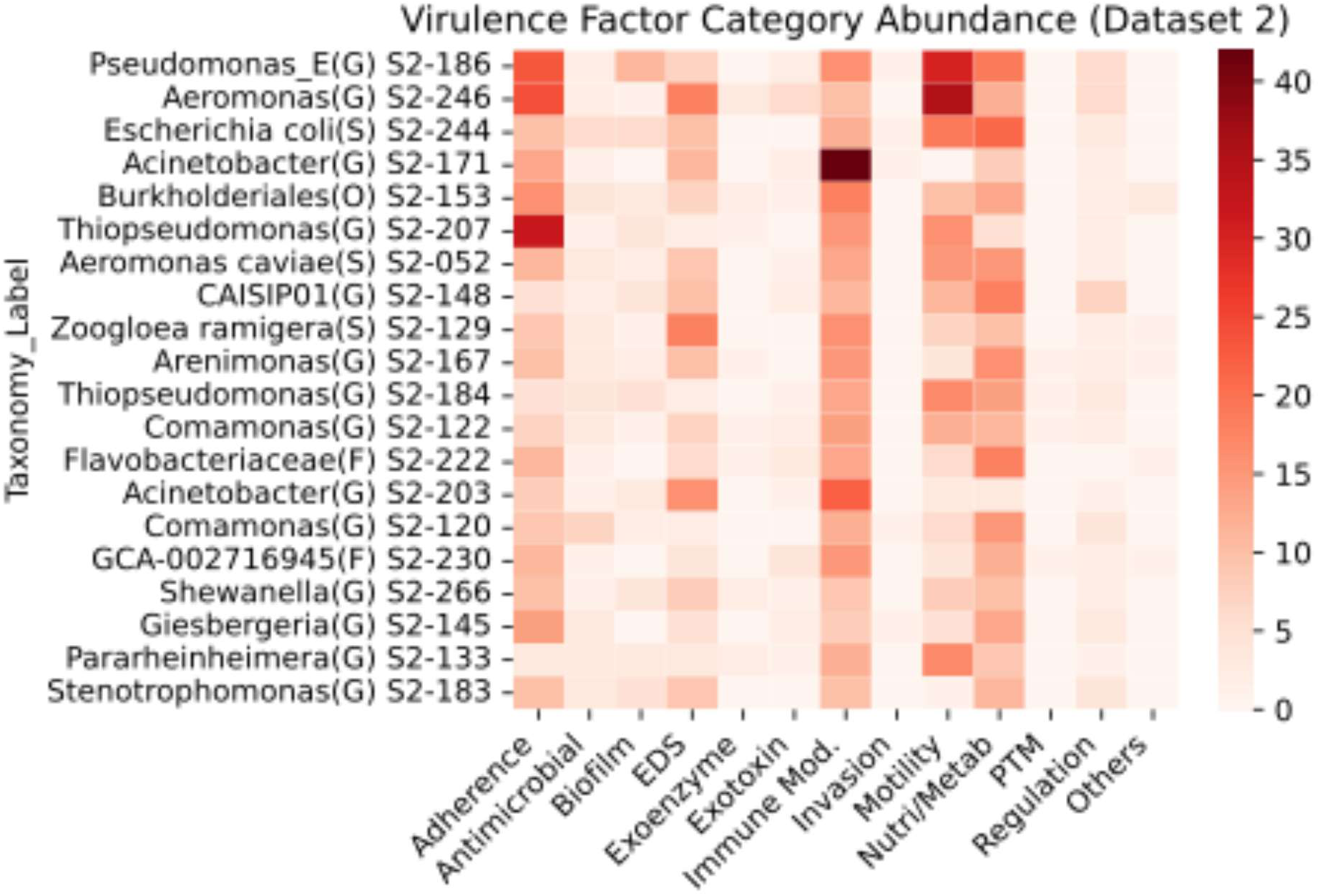
Heatmap showing virulence factor (VF) counts per VF category for the top 20 MAGs containing the most overall VFs in Dataset 2. VF categories on the X-axis were abbreviated (e.g., EDS = Effector Delivery System; Immune Mod. = Immune Modulation; Nutri/Metab = Nutritional/Metabolic Factor; PTM = Post-Translational Modification). The Y-axis lists MAGs by their most resolved taxonomic rank and numerical identifier, where S2 indicates the dataset to which the study belongs (Shahar et al., 2024a), and the numbers following it indicate the bin number assigned by Maxbin2 v2.2.7. Color intensity reflects VF count, with darker colors indicating higher counts. Taxonomic abbreviations: O = Order, F = Family, G = Genus, and S = Species. Supplementary Table 2 contains the complete table for all MAGs in Dataset 2.

### NMDS Analysis of ARG Read Counts Across Two Datasets

Figure 6 shows an NMDS plot using the ARG abundances for Dataset 1. Samples 2, 11, and 22 from Yogurt Canyon are separated from the main cluster, indicating distinct ARG profiles in these samples. In the NMDS ordination plot for the ARG abundances for Dataset 2 (Figure 7), samples from Radio Club generally clustered together but exhibited some temporal shifts, whereas Boca Rio samples were more dispersed, reflecting higher variability in ARG profiles.

**Figure 6.**
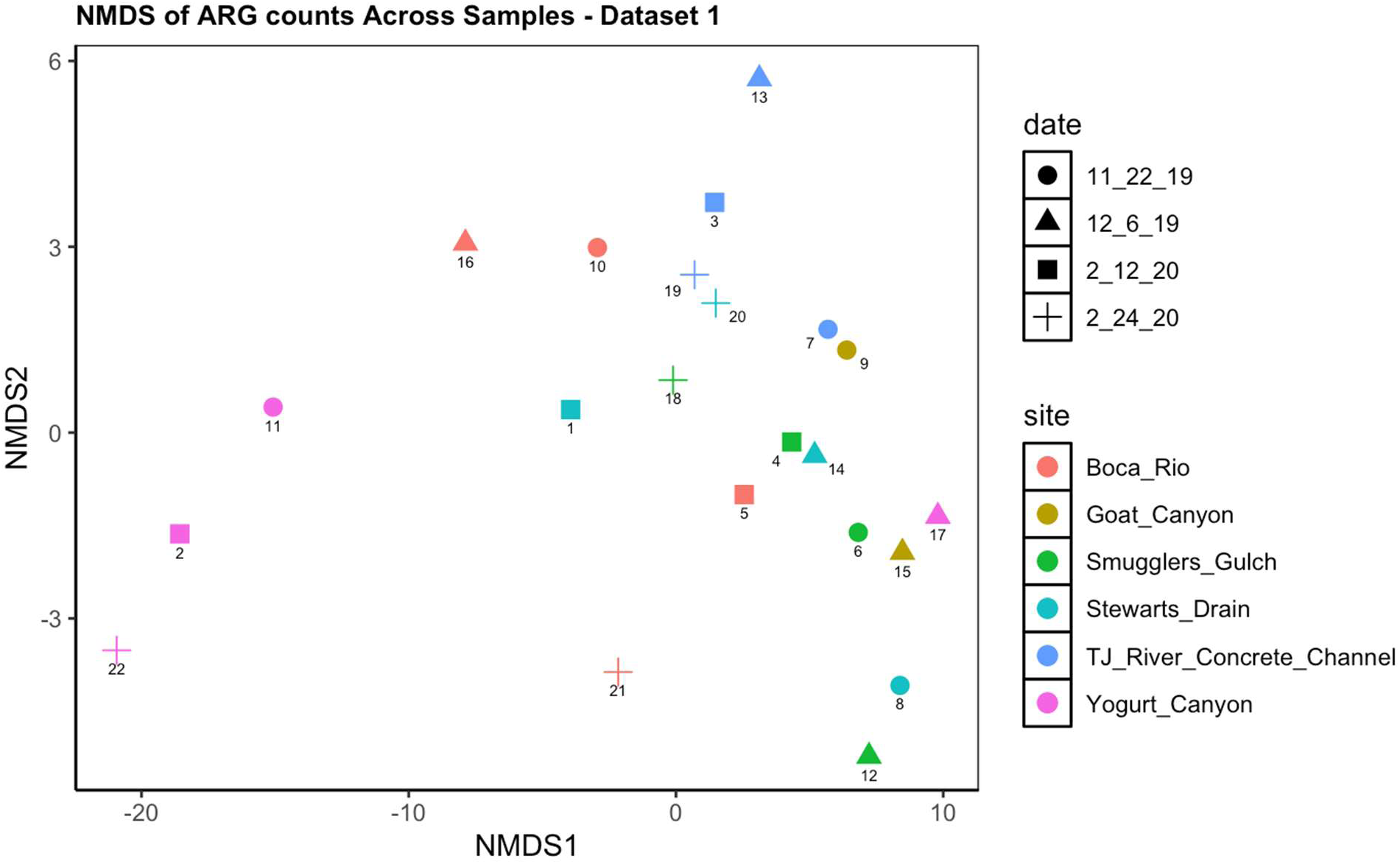
NMDS plot of ARG read counts across samples for Dataset 1. Read counts were robust centered log-ratio (rCLR) transformed, and Euclidean distances were calculated to quantify dissimilarities between samples prior to NMDS. Each point represents an individual sample, with colors indicating sampling site and shapes indicating sampling date. Numbers next to points correspond to individual samples. NMDS stress = 0.067

**Figure 7.**
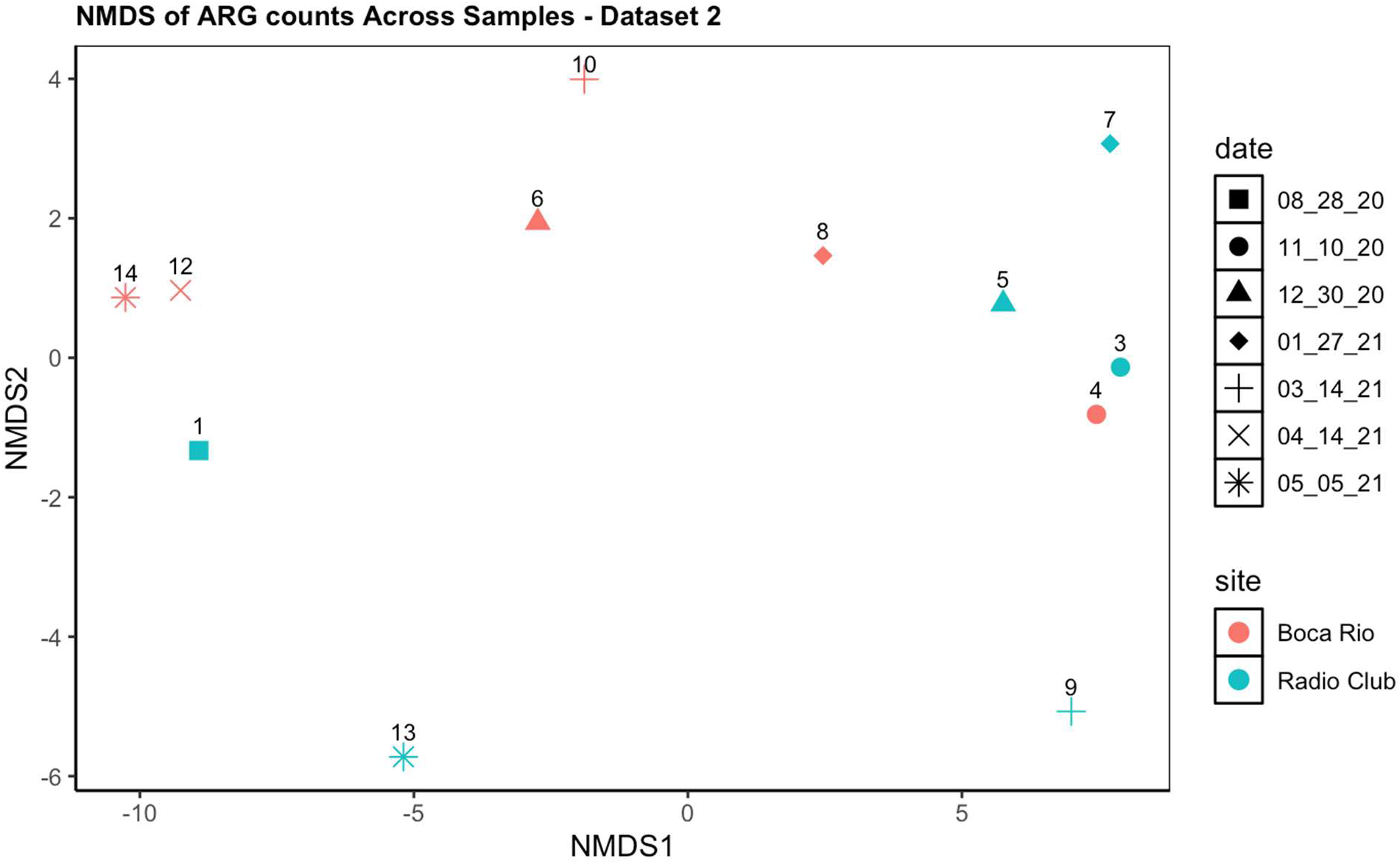
NMDS plot of ARG read counts across samples for Dataset 2. Read counts were robust centered log-ratio (rCLR) transformed, and Euclidean distances were calculated to quantify dissimilarities between samples prior to NMDS. Each point represents an individual sample, with colors indicating sampling site and shapes indicating sampling date. Numbers next to points correspond to individual samples. NMDS stress = 0.042

Consistent with the ordination patterns, the PERMANOVA results showed a significant effect of site for Dataset 1 (pseudo-F = 1.9; R² = 0.37; p = 0.02), whereas no significant site-level differences were detected for Dataset 2. Sampling date had no significant effect in either dataset. Overall, the weak clustering patterns observed in the NMDS plots indicate that ARG profiles are heterogeneous and not strongly structured by site or sampling date, particularly in Dataset 2.

### Antibiotic Resistance Genes Across Samples

The top 10 ARGs were identified based on total read counts derived from unbinned contigs. For each ARG, the total read count was calculated by summing its read counts across all samples. The top 10 ARGs with the highest total read counts are shown in Table 1 and Table 2 for Datasets 1 and 2, respectively. In Table 1, tet(39), an efflux-associated gene, had the highest total read count (5,255 reads) followed by tet(C) with 4617 reads. Both inactivation- and efflux-associated genes were represented among the top 10 ARGs, with inactivation genes being more numerous than efflux genes.

**Table 1.** List of Top 10 ARGs for Dataset 1.

| ARG | ARG Category | Total Read Count |
| --- | --- | --- |
| tet(39) | Efflux | 5255 |
| tet(C) | Efflux | 4617 |
| OXA-1243 | Inactivation | 3771 |
| aadA25 | Inactivation | 3736 |
| ANT(3'')-II-AAC(6')-IId | Inactivation | 2932 |
| tet(E) | Efflux | 2871 |
| qacEdelta1 | Efflux | 2385 |
| APH(6)-Id | Inactivation | 2357 |
| CfxA5 | Inactivation | 1502 |
| aadA2 | Inactivation | 1234 |

**Table 2.** List of Top 10 ARGs for Dataset 2.

| ARG | ARG Category | Total Read Count |
| --- | --- | --- |
| qacEdelta1 | Efflux | 682 |
| cmlA8 | Efflux | 591 |
| OXA-1163 | Inactivation | 260 |
| Mef(En2) | Efflux | 248 |
| MexF | Efflux | 230 |
| acrB | Efflux | 220 |
| AcrF | Efflux | 191 |
| MexK | Efflux | 175 |
| mdtF | Efflux | 157 |
| mdtC | Efflux | 142 |

In Table 2, qacEdelta1 (efflux gene) had the highest total read count (682 reads), followed by cmlA8 (591 reads). Most of the top ARGs were efflux-associated genes, with only OXA-1163 classified as an inactivation gene. Overall, efflux-associated ARGs dominated the top 10 in Dataset 2, indicating that efflux mechanisms were the most frequently detected ARG types in this dataset.

### Network Analysis of ARG-MAG Correlations

Figures 8 show correlation networks between MAGs and ARGs for Datasets 1 (Figure 8A) and 2 (Figure 8), respectively. Only positive correlations were considered, as these reflect co-occurrence patterns in which higher abundance of a MAG is associated with higher counts of specific ARGs. Overall, both networks exhibited strong positive correlations, with Spearman correlation coefficients of r ≥ 0.90 for Dataset 1 and r ≥ 0.95 for Dataset 2. MAGs are labeled at their lowest resolved taxonomic level (domain, family, genus, or species) and the numbers next to it correspond to the bin numbers assigned by Maxbin2 v2.2.7.

**Figure 8.**
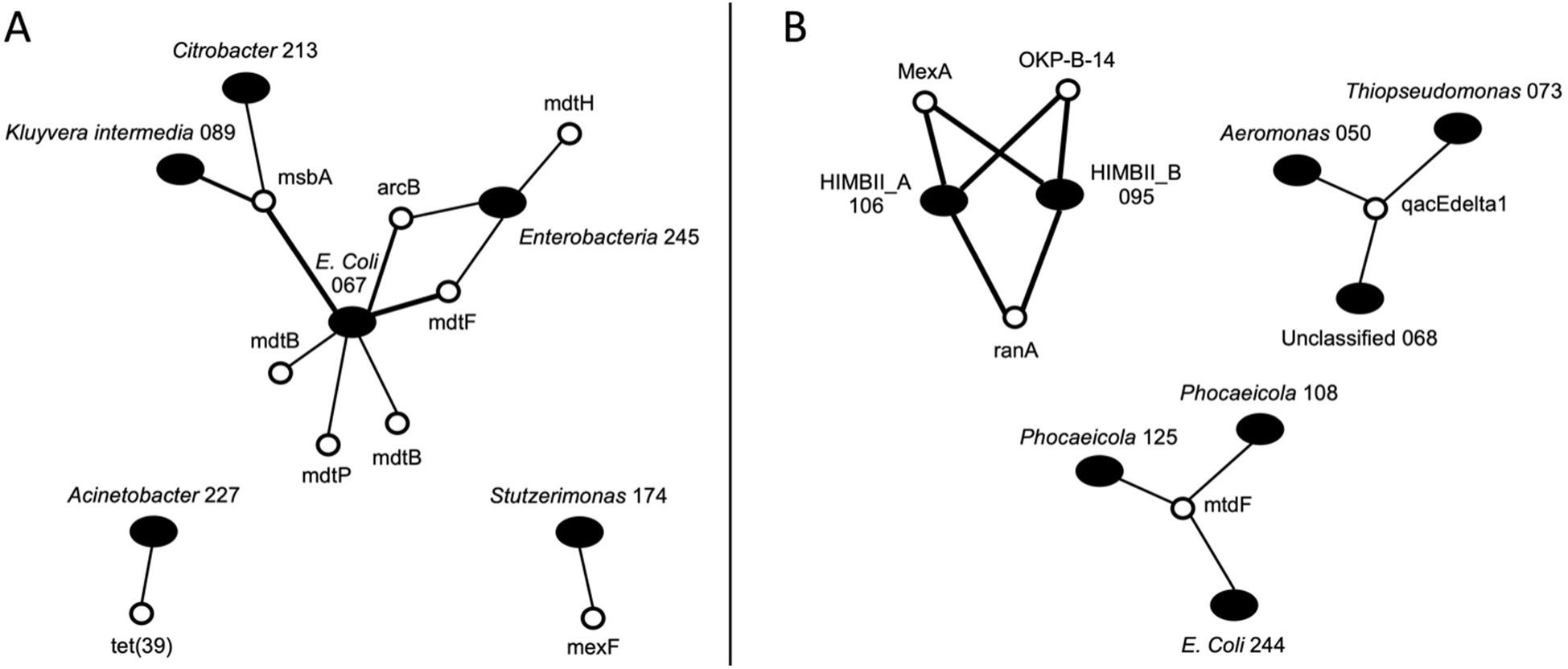
Positive network for ARGs and MAGs for (A) Dataset 1 and (B) Dataset 2. The filled ovals (nodes) indicate bacterial MAGs, and the open circles indicate ARGs, with the lines (edges) represent significant positive correlations (Spearman rank) between MAGs and ARGs. All correlation coefficients ranged from 0.9 to 1.0 (perfect correlation) with the thickness of the lines indicate the strength of the correlation. Clusters of connected nodes indicate groups of MAGs and ARGs that are highly correlated across samples. MAGs are labeled at their most resolved taxonomic rank, and the numbers correspond to the bin numbers assigned by MaxBin2 v2.2.7.

In Dataset 1 (Figure 8A), the network involving *E. coli* (S) 067 exhibited multiple positive correlations with efflux ARGs, including mdtE, mdtP, mdtB, mdtF, acrB, and msbA (Nishino et al., 2008; Wright et al., 2025). In addition, msbA was positively correlated with *Kluyvera intermedia_A* (S) 089 and *Citrobacter* (G) 213, indicating shared ARG-MAG associations. A family-level MAG, *Enterobacteriaceae* (F) 245, also showed positive correlations with efflux-associated genes (acrB, mdtH, and mdtF). In contrast, *Acinetobacter* (G) 227 was associated with a single efflux ARG (tet(39)), which ranked first among the top 10 ARGs in Dataset1 (Agersø & Guardabassi, 2005). Similarly, *Stutzerimonas* (G) 174 exhibited a single positive correlation with mexF, an efflux ARG, indicating limited ARG connectivity, consistent with previous reports describing efflux-mediated resistance mechanisms in *Stutzerimonas* species (Shi et al., 2024).

In Dataset 2 (Figure 8B), the positive correlation network resolved into three distinct clusters. One cluster included efflux genes MexA (Shi et al., 2024) and RanA (Yang et al., 2024), as well as the inactivation gene (OKP-B-14), (Melano et al., 2006), all of which were positively correlated to HIMB11 MAGs. A second cluster included qacEdelta1, an efflux ARG (Jiang et al., 2025) that ranked first among the top 10 ARGs in Dataset 2, and was positively correlated with MAGs assigned to *Aeromonas*, *Thiopseudomonas*, and unclassified bacteria, reflecting a broader taxonomic distribution. A third cluster consisted of the efflux gene mdtF, which also ranked among the top 10 ARGs in Dataset2 and was positively correlated with *E. coli* and *Phocaeicola* MAGs.

## Discussion

This study aimed to develop and evaluate a reproducible co-assembly-based metagenomic workflow to reconstruct microbial genomes and associate virulence factors with MAGs, while also identifying antimicrobial resistance genes (ARGs) in the unbinned fraction of co-assembled contigs across environmental samples from the Tijuana River. By integrating co-assembly, binning, functional annotation, and correlation network analyses, this workflow provides a framework for examining microbial community structure alongside resistance and virulence potential.

Following quality control, a co-assembly-based metagenomic workflow was adopted to recover longer contigs and improve gene-level resolution compared to read-based classification approaches such as Kaiju. Unlike read-based methods, which assign taxonomy and function directly to short reads, co-assembly enables the grouping of reads into contiguous sequences, preserving gene order and genomic context. This is particularly advantageous for downstream genome reconstruction and functional analysis (Erkorkmaz et al., 2025). Consequently, co-assembly was the most computationally demanding step in the workflow, requiring high-memory servers for both datasets (Lynn & Gordon, 2025). Despite these resource requirements, co-assembly enabled the recovery of metagenome-assembled genomes (MAGs) of varying completeness, thereby increasing confidence in downstream functional analyses. Separating binned MAGs from unbinned contigs further improved biological interpretability, as the unbinned fraction likely represents mobile genetic elements, whereas MAGs reflect more stable genomic content (Abramova et al., 2024).

The impact of this separation was evident in the distribution and characteristics of binned and unbinned sequences across both datasets. Although individual MAGs were substantially longer than unbinned contigs, the highly fragmented unbinned fraction collectively contributed a greater total sequence length in both datasets, suggesting that these sequences largely derived from low-abundance organisms, repetitive regions, or mobile genetic elements that remain unassigned during binning (Maguire et al., 2020). While a subset of bins reached high completeness (>90%), most MAGs were of low to moderate completeness, reflecting the inherent challenges of co-assembly and binning in complex environmental metagenomes, particularly for low-abundance taxa and fragmented genomic regions (Zhou et al., 2022).

Despite the wide range of genome sizes and variable completeness observed across recovered MAGs, the co-assembled genome set was sufficient to resolve meaningful community-level patterns. NMDS ordination of MAG relative abundance for Dataset 1 showed that most samples clustered closely together, indicating broadly similar microbial community compositions across sites (Figure 2). However, Yogurt Canyon and Boca Rio samples were separated from the main cluster, suggesting distinct site-specific MAG compositions. Boca Rio is on the coast where the river mixes with marine waters, while Yogurt Canyon is close to the border where the sewage spills over regularly. A similar pattern was reported by (Allsing et al., 2022) based on short-read taxonomic analysis using Kaiju of the same data. PERMANOVA analysis confirmed that sampling site significantly influenced MAG composition, whereas collection date did not, indicating that spatial environmental factors exert a stronger influence than temporal variation. Similar trends were observed in Dataset 2. Radio Club (near border) samples clustered tightly together, reflecting high within-site similarity, while a subset of Boca Rio (mixed river and coastal marine) samples formed a distinct cluster in ordination space (Figure 2). PERMANOVA again identified site as a significant determinant of MAG composition, with no detectable effect of collection date consistent with the spatial structuring of microbial communities determined with the same dataset based on short-read Kaiju analysis (Shahar et al., 2024a).

While genome-resolved analyses revealed clear site-specific structuring of microbial communities, examining ARG profiles within the unbinned fraction provides additional insight into functional traits that may not be captured within MAGs. For Dataset 1, NMDS ordination of ARG relative abundance showed that most Yogurt Canyon samples were separated from the main cluster, indicating distinct ARG compositions relative to other sites (Figure 6). This spatial pattern was supported by PERMANOVA, which found sampling site to be a significant factor, but not collection data. In contrast, ARG ordination for Dataset 2 did not reveal clear clustering by site (Figure 7), and there was no significant association with ARG composition. This lack of spatial structure in ARG profiles may reflect the more dynamic nature of resistance genes, which are often associated with mobile genetic elements (Maguire et al., 2020).

While ARG profiles highlight resistance potential within the community, virulence factors provide complementary insight into microbial strategies for host interaction and environmental adaptation. To explore this functional potential, we examined MAGs for predicted virulence factors, which offer insight into how microbes may interact with hosts or adapt to their environments. MAGs from Dataset 1 showed substantial variation in VF profiles, with Immune Modulation, Motility, and Nutritional/Metabolic factors being the most prevalent (Figure 4). Notably, *Pseudomonas_E* MAGs were enriched in motility and immune modulation virulence factors. Members of this genus, including species such as *Pseudomonas aeruginosa*, which are opportunistic pathogens, are known to use flagella and pili for motility and possess mechanisms to modulate host immune responses (Sultan et al., 2021). MAGs within the class Campylobacteria displayed high immune modulation virulence factors. Members of this class include pathogenic *Campylobacter* species, which are well-known foodborne enteric pathogens whose virulence traits can influence host immune responses during gastrointestinal infection (Bolton, 2015). MAGs classified to the genus *Aliarcobacter* were enriched in immune modulation virulence factors. Members of this genus have been identified as emerging opportunistic and potentially zoonotic human pathogens, with comparative genomic analyses revealing virulence determinants linked to host interaction, invasion, and immune modulation, including immune evasion mechanisms (Chuan et al., 2022).

MAGs from Dataset 2 also exhibited variable VF profiles, with Adherence, Immune Modulation, Motility, and Nutritional/Metabolic factors being common (Figure 5). In this dataset, *Pseudomonas_E* and *Aeromona* MAGs were enriched in Motility VFs, a MAG from the genus *Acinetobacter* showed high Immune Modulation VFs, and a *Thiopseudomonas* MAG displayed elevated Adherence VFs. *Aeromonas* species are primarily water and foodborne bacteria that can cause gastrointestinal and extraintestinal infections in humans, particularly in immunocompromised individuals (Tomás, 2012). Members of the genus *Acinetobacter* are opportunistic human pathogens whose virulence factors have a role in modulating host immune responses (Shadan et al., 2023).

The vast majority of the ARGs were detected in the unbinned fraction of both datasets, supporting the notion that the ARGs were plasmid associated. To assess which resistance genes prevail across both datasets, we performed BLASTp on the unbinned fraction of contigs against the CARD database, focusing on both inactivation- and efflux-associated genes. ARG read counts were substantially higher in Dataset 1, reaching the thousands, compared to the hundreds observed in Dataset 2. Additionally, the top 10 ARGs in each dataset did not overlap, highlighting pronounced differences in resistome composition between the two datasets.

In Dataset 1, both efflux and inactivation associated ARGs were prevalent among the top-ranked genes, with inactivation genes occurring more frequently overall (Table 1). The high abundance of tetracycline resistance genes, including tet(39) and tet(C), suggests sustained selective pressure for tetracycline resistance and underscores the importance of active efflux as a primary resistance mechanism in this environment. Tet (39) is a tetracycline efflux pump found in Gram-negative bacteria, including *Brevundimonas*, *Stenotrophomonas*, *Enterobacter*, *Alcaligenes*, *Acinetobacter*, and *Providencia*. It is often plasmid-associated, facilitating horizontal transfer across bacterial species. Tet (C) is a tetracycline efflux pump found in many species of Gram-negative bacteria. It is typically found in plasmid DNA (Alcock et al., 2023b). Similar ARG classes, including efflux-associated (tet, qacEdelta1) and inactivation-associated (aadA, OXA) genes, have been reported as dominant resistance determinants in previous metagenomic studies of the Tijuana River watershed (Allsing et al., 2022).

In contrast, Dataset 2 was dominated by efflux-associated ARGs, with qacEdelta1 and cmlA8 among the most abundant (Table 2). The qacEdelta1gene is an efflux resistance gene that confers resistance to antiseptics and disinfectants and is commonly associated with plasmids, while cmlA8 is a chloramphenicol resistance gene that functions through an efflux mechanism (Alcock et al., 2023b). Similar to previous studies, Dataset 2 was dominated by efflux-associated ARGs, including qacEdelta1, chloramphenicol exporters, and OXA-type β -lactamases, indicating consistent resistance mechanisms across samples (Shahar et al., 2024b).

Because the ARGs did not belong to genome bins and were likely on plasmids we attempted to identify potential ARG hosts by creating correlation networks between MAG and ARG abundances. In both datasets, we found highly positive ARG-MAG correlations (rho ≥ 0.9), strongly indicating bacteria that could be the potential plasmid carriers (Figure 8). The highly correlated efflux-associated ARGs in Dataset 1 were associated with *Enterobacteriaceae*, *E. coli*, *Kluyvera intermedia_A*, and *Citrobacter* MAGs (Figure 8A). Notably, *E. coli* MAGs exhibited multiple positive correlations with efflux-related ARGs, including mdtE, mdtP, mdtB, mdtF, acrB, and msbA, consistent with well-documented efflux-mediated resistance mechanisms in enteric bacteria. In contrast, *Acinetobacter* showed an association with the efflux-associated gene tet(39), while *Stutzerimonas* displayed a strong positive correlation with the mexF efflux pump.

In Dataset 2, the efflux genes mexA and ranA and the inactivation gene OKP-B-14, were strongly correlated with two unknown species of Rhodobacteraceae (Figure 8B). A second cluster centered on qacEdelta1, an efflux-associated ARG, which was positively correlated with MAGs assigned to *Aeromonas*, *Thiopseudomonas*, and one unclassified bacterial MAG. A third cluster consisted of the efflux gene mdtF, which showed positive correlations with *E. coli* and *Phocaeicola* MAGs. These ARG-MAG associations in Dataset 2 possibly reflect horizontal transmission of plamids among unrelated bacteria.

## Conclusion

A key strength of the proposed workflow is its flexible design. This design enabled the same workflow to be applied to both datasets with minimal modification, demonstrating its robustness and general applicability to similar metagenomic datasets. Despite these advantages, the workflow has certain limitations. Co-assembly, binning, and BLAST-based functional annotation are computationally demanding and may not be feasible for extremely large datasets without access to high-performance computing infrastructure. In addition, the accuracy of functional annotation depends on the quality and coverage of available reference databases, which may limit the detection of novel or poorly characterized genes. Future improvements could include incorporating new binning algorithms or combining the outputs of multiple independent binning approaches, using seed co-assemblies to allow for the inclusion of more metagenomes, and the incorporation of long-read sequencing to enhance genome completion. Our general workflow framework should make such improvements easy to enable.

## Data availability

The metagenomic datasets used for testing are publicly available in the European Nucleotide Archive, project accession number PRJEB57859 and at the NCBI Sequence Read Archive (SRA), Bioproject PRJNA1423341. The complete pipeline with all details on software and database requirements as well as space needs can be found at https://github.com/ddkrutkin/ARGus/tree/main. The statistical analysis code used in this article is available at https://github.com/nivepulsub/TJ_River_Analysis/tree/main. Supplementary material has been made available at Zenodo DOI: <u>10.5281/zenodo.19193950</u> https://zenodo.org/records/19193951.

## Acknowledgments

We thank Dr. Arun Sethuraman and Dr. Luke Miller for helpful comments on the manuscript and Alison Peterson for critical infrastructure support.

